# Microengineered three-dimensional collagen fiber landscapes with independently tunable anisotropy and directionality

**DOI:** 10.1101/2020.06.12.148346

**Authors:** Adeel Ahmed, Indranil M. Joshi, Mehran Mansouri, Stephen Larson, Shayan Gholizadeh, Zahra Allahyari, Farzad Forouzandeh, David A. Borkholder, Thomas R. Gaborski, Vinay V. Abhyankar

## Abstract

Fibrillar collagens are structural proteins in the extracellular matrix (ECM), and cellular processes, including differentiation, proliferation, and migration, have been linked to the orientation (directionality) and alignment (anisotropy) of collagen fibers. Given the importance of cell-substrate interactions in driving biological functions, several microfluidic approaches have demonstrated three-dimensional (3D) collagen gels with defined fiber properties that enable quantitative correlations between structural cues and observed cell responses. Although existing methods provide excellent definition over collagen fiber anisotropy, independent control over both anisotropy and directionality (that we collectively refer to as the collagen landscape) has not been demonstrated. Therefore, to advance collagen microengineering capabilities, we present a user-friendly approach that uses controlled fluid flows within a non-uniform microfluidic channel network to create well-defined collagen landscapes. We demonstrate capabilities including i) control over fiber anisotropy, ii) spatial gradients in fiber anisotropy, iii) defined fiber directionality, and iv) multi-material interfaces. We then show that cells respond to the microengineered topographic cues by aligning along the anisotropy domains and following fiber directionality. Finally, this platform’s modular capability is demonstrated by integrating an ultrathin porous parylene (UPP) membrane on the microengineered collagen as a mask to control cell-substrate interactions.

## 1. INTRODUCTION

In native tissue, the extracellular matrix (ECM) is a heterogeneous macromolecular network that is dynamically remodeled in response to biomechanical and biochemical stimuli^1^. These interactions induce the local reorganization of type 1 collagen (COL1) fibers into aligned (anisotropic) fiber domains with defined orientation (directionality)^2,3^. Here, we refer to fiber anisotropy and directionality as components of the overall collagen landscape, which is known to influence the motility of T-cells^4^, guide endothelial cell alignment^5^, regulate cell differentiation^6^, and direct the wound healing process^7^. In human biopsy samples, characteristics of the COL1 landscape at the tumor-stroma boundary accurately predict the long-term survival for breast and pancreatic cancer patients^8,9^. Given the importance of COL1 fiber properties in a broad range of biological processes, there have been significant *in vitro* efforts to replicate the characteristics of collagen landscapes to study cellular responses in a controlled manner^10^.

Microfluidic systems are particularly well-suited for advanced cell culture applications owing to the precise control over fluid flows, soluble factors, and cell patterning capabilities made possible by favorable scaling effects^11,12^. The formation of anisotropic COL1 fibers has been demonstrated in microscale systems by manipulating neutralized collagen solutions during the self-assembly process by controlling the motion of magnetic beads^13,14^, introducing shear flows^15–17^, and applying mechanical strain^18^. Alternatively, fibroblast-based remodeling and decellularization of pre-gelled collagen have achieved highly anisotropic collagen environments to study directed cell motility^19,20^. Although current microfluidic and cell-based collagen fabrication techniques offer considerable definition over fiber anisotropy, they have not yet demonstrated independent control over fiber anisotropy *and* directionally (e.g., turns and curves) to replicate the *in vivo* collagen landscape.

Over the past ten years, extrusion-based bio-fabrication techniques have matured and can now achieve increasingly complex shapes with precise spatial control over the extruded strands^5,21^. The number of available printing materials has also expanded, and collagen-blended bio-inks and sacrificial layers are commonly used as construction materials^21–23^. Although extrusion-based approaches accurately control strand position and directionality, they cannot achieve the internal collagen fiber anisotropy that is possible using microengineering and cell-based methods. To address the need for *in vitro* systems to study cell responses on well-defined collagen landscapes, we present an approach that combines microengineered fiber anisotropy with controlled fiber directionality in a single system.

This work introduces a microfluidic platform that uses a multi-port channel network with non-uniform channel dimensions to impose controlled extensional strain on neutralized COL1 solutions to control fiber directionality and promote anisotropy. First, we demonstrate: i) defined anisotropic collagen fiber domains on the millimeter scale, ii) spatial gradients in fiber anisotropy, iii) control over fiber directionality, and iv) multi-material interfaces with controlled anisotropy. Second, we address the practical challenge of introducing cells into microfluidic systems and show that single cells and multi-cellular sheets respond to the topography presented by our microengineered collagen landscapes. Finally, we present the modular capability of our platform by incorporating a microporous, ultrathin porous parylene (UPP) membrane^24^ on top of the collagen landscape as a method to control the degree of interaction between the cells and the underlying fiber structures.

## 2. MATERIALS AND METHODS

### Soft Lithography

Polydimethylsiloxane (PDMS) microfluidic channels were fabricated using standard soft lithography techniques^25^. Briefly, photoresist (SU-8 3050, Kayaku Advanced Materials, Massachusetts, USA) was spin-coated to a thickness of 60μm on a 4” diameter silicon wafer (University Wafers, South Boston, USA), followed by a soft bake at 95°C for 30 minutes. The photoresist was then exposed to UV (250mJ/cm^2^) through a high-resolution printed photomask, baked for 45 minutes at 90°C, and then developed in SU-8 developer for 20 minutes. The wafer was then rinsed with IPA and dried under an air stream prior to storage. A 10:1 pre-polymer base to crosslinker ratio (Sylgard 184, Dow Corning, Midland USA) was mixed and then degassed in a vacuum chamber for 30 minutes until a bubble-free solution was achieved. The PDMS solution was then poured over the SU-8 mold and cured for 2 hours at 80°C on a hotplate. The resulting PDMS channels were then gently removed from the wafer and cut to size with a razor blade while fluidic access ports were cored with a 1 mm biopsy punch (World Precision Instruments, USA). PDMS channels were then cleaned using 70% ethanol, followed by a DI water rinse and then stored covered until use. Channel dimensions are shown in Figure S1, and the total volume is ~ 5 μL.

### PDMS passivation and glass functionalization

Molded PDMS microchannels were sonicated in 70% ethanol for 5 minutes, followed by a sterile DI water rinse, and dried in a biosafety cabinet. The channels were then immersed in a 1% (w/v) solution of bovine serum albumin (BSA) in sterile PBS (Fisher Scientific, USA) for 4 hours at 4°C to passivate the surfaces. The BSA solution was then aspirated, rinsed with PBS, and the channels were allowed to dry in a biosafety cabinet. BSA-passivated channels were stored at 4°C for up to 2 days before use.

Glass coverslips (25×50mm, #1.5, Globe Scientific, USA) were sonicated in 70% ethanol, dried in air, and then exposed to O_2_ plasma for 1 minute at 600mTorr using the high-power setting (Harrick Plasma, Ithaca, NY, USA). The coverslips were then submerged in a solution containing 2% (v/v) aminopropyltriethoxysilane (APTES) (Millipore Sigma, Burlington, USA) with 90% (v/v) ethanol, 5% (v/v) deionized water and 3% (v/v) glacial acetic acid. The coverslips were placed on a rocker plate for 5 minutes, rinsed with ethanol, and baked on a hot plate at 110°C for 10 minutes. Next, a 0.08% (w/v) solution of poly(octadecene maleic alt-1-anhydride) (POMA) (Millipore Sigma, Burlington, USA) was dissolved in tetrahydrofuran (Millipore Sigma, Burlington, USA) and spin-coated on to the APTES functionalized coverslips at 1000 rpm for 20 seconds. The slides were then baked at 120°C for 1 hour and stored at room temperature until use. This process resulted in a surface coating of POMA to promote covalent attachment of collagen to the surface during gelation.

### Collagen neutralization

Type I bovine atelocollagen (COL1) (Nutragen, Advanced Biomatrix, San Diego, USA) (417 μl) was diluted to 2.5 mg ml^−1^ in 10X PBS (100μl) (Alfa Aesar, Ward Hill, USA) and ultrapure water (478μl) and then neutralized using 1M NaOH (5.4μl) (VWR, Randor, USA) to a pH of 7.5 immediately before use. A 2.5 mg ml^−1^ collagen solution containing 0.48% (v/v) hyaluronic acid (HA) (Fisher Scientific, AAJ6699303, MW > 1MDa) was created by replacing the ultrapure water with an equivalent volume of a 10mg ml^−1^ HA solution in PBS.

### Collagen Rheometry

Rheometry was carried using a hybrid rheometer with a temperature-controlled stage (TA instruments HR-2, Delaware, USA). A 40mm truncated cone geometry was used (TA instruments part no: 113847) with a sample volume of 300μl and a gap of 25μm. A neutralized collagen solution was prepared before loading as described, and the rheometer temperature was maintained at 21°C. The complex viscosity was measured under 10% strain at 1Hz for 10 minutes. The complex viscosity used was measured at 300s after the start of the experiment, representing the time between collagen preparation and injection into the channels. Results are shown in the supplement (Figure S2)

### Collagen injection process

PDMS microfluidic channels were reversibly sealed against POMA-functionalized glass coverslips via conformal contact. A neutralized 2.5 mg ml^−1^ solution of collagen was hand injected into the microfluidic channel network using a pipette inserted into port 3 with all the other ports open to the atmosphere (see Figure 1). The pre-filling step ensured the bubble-free collagen introduction during the second injection step. Following pre-filling, ports 3 and 5 were sealed using adhesive tape (Scotch brand, 3M, USA), and a second collagen solution (COL1) or collagen-HA (COL1-HA) was injected at a constant flow rate of 500 μl min^−1^ from port 2 using a syringe pump (NE-4000, New Era Pump Systems Inc, USA). Priming and injection steps were carried out with the microfluidic chips at 21°C and with the collagen stored on ice. Samples were then transferred to a 37°C incubator for gelation.

**Figure 1:**
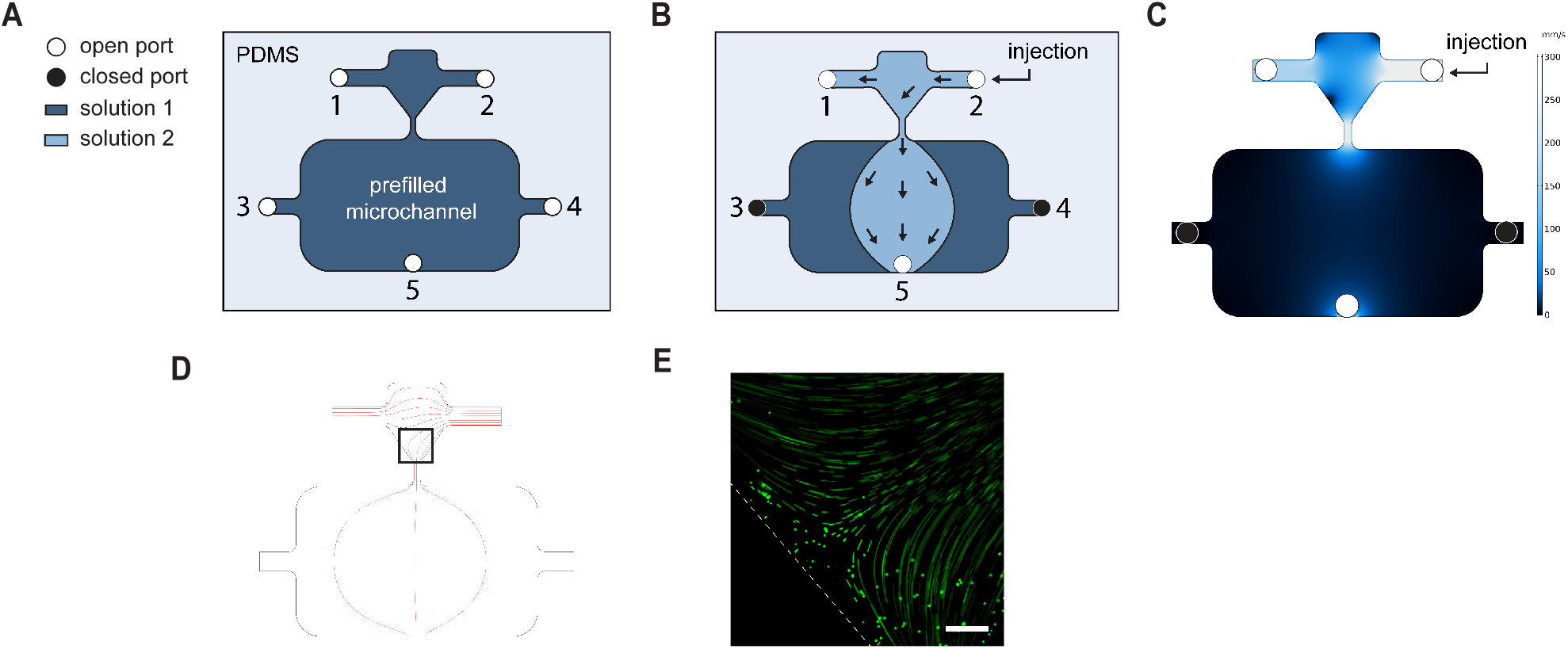
**(A)** Top-down view of the PDMS channel network. Open access ports are shown as white circles and numbered. The ports are open during pre-filling (priming) with collagen (dark blue) from port 3. **(B)** Solid back circles indicate closed ports. An injection of a second collagen solution (light blue) was performed Q = 500*μ*L min^−1^ along the path 2-1 and 2-5 to impose an extensional strain on the collagen solution. **(C)** A COMSOL simulation shows the velocity along the flow path (2-1 and 2-5) with lighter shades of blue indicating a higher magnitude. **(D)** Fluid flow streamlines simulated using COMSOL with the black box depicting the ROI for panel E. **(E)** Streamline image of fluorescent microbeads flowing from ports 2-5 shows that the collagen flow path was consistent with the simulated results. Scale bar = 50*μ*m.

5μm streptavidin functionalized paramagnetic beads (Part no: UMC0101, Bangs Laboratories, Indiana, USA) were conjugated to Atto-500 Biotin (Sigma Aldrich, USA) and were incorporated into the COL1/HA solution at a concentration of 0.1 mg ml^−1^ and used to identify the interface between the COL1 and COL1/HA regions. The solution was then injected as described above, and fluorescence imaging was carried out using an Olympus IX-81 inverted microscope using the appropriate filter set. Contrast adjustment was carried out to enhance visual clarity using ImageJ (National Institutes of Health, USA)

### Flow Simulation

A steady-state 3D simulation was performed using laminar flow physics of COMSOL Multiphysics®. The 2D geometry was designed in Adobe Illustrator (Adobe Systems, California, USA) and imported into COMSOL and a 60μm thickness was applied to the geometry. A fully developed flow rate of 500 μl min^−1^ inlet boundary condition was applied to port 2, while the outlet boundary condition of pressure *P* = 0 was applied to ports 1 and 4. Wall boundary conditions were applied to all other boundaries. A combination of tetrahedral, prisms, and triangle mesh elements were applied to the geometry, and a mesh independency study was carried out with stable results at ~1,170,000 elements. Steady-state velocity magnitude contour was mapped on two lines from ports 2-1 and in the constriction from ports 2-5. The extensional strain rate was calculated by determining the slope of the line plot at the desired coordinates.

### Cell Culture

MDA-MB-231 cells (ATCC, Manassas, USA) were cultured in Dulbecco’s modified eagle medium (Gibco, Grand Island, USA) with 10% fetal bovine serum (Gibco, Grand Island, USA) at 37°C with 5% CO_2_. Cells were used between passages 5 and 7. The media was changed every 48 hours. Human Umbilical Vein Endothelial Cells (HUVEC) (Thermo Fisher Scientific, USA) were cultured in medium M200 (Thermo Fisher Scientific, USA) supplemented with a low serum growth supplement kit (Thermo Fisher Scientific, USA). HUVECs were used between passage 1-3 Cells were enzymatically dissociated using TrypLE (ThermoFisher Scientific, MA, USA) for 5-7 minutes and centrifuged at 250G for 5 minutes, and resuspended. MDA-MB-231 cells were seeded at a density of 30,000 cm^−2^, and HUVECs were seeded at 20,000 cm^−2^. Before seeding, PDMS channels were lifted off from the POMA functionalized coverslips to enable direct access to the collagen gel. A 2mm × 20mm^2^ laser-cut poly(methyl methacrylate) (PMMA) well was cleaned with ethanol and attached to the region surrounding the gel using a pressure-sensitive adhesive (MP468, 3M, USA).

### Immunostaining and imaging

Twenty-four hours after seeding, cells were fixed in 3.7% paraformaldehyde in PBS for 15 mins. Cells were then permeabilized in Triton X-100 (0.1%) for 10 mins and washed with PBS Tween-20 (PBST). Cells were blocked in 40 mg ml^−1^ BSA (Alfa Aesar, Ward Hill, USA) for 30 minutes at room temperature. Cells were labeled with Hoechst 33342 (300 nM) (Molecular Probes, USA) for 10 min and AlexaFluor 488 conjugated phalloidin (ThermoFisher Scientific, MA, USA) (1:400) for 15 mins to visualized nuclei and actin fibers, respectively. Finally, cells were washed with PBS tween-20 and stored in PBS. Fluorescence imaging was performed an Olympus IX-81 inverted microscope with CellSens software, and all image collection settings were consistent across experimental sets to allow comparison. Image processing was carried out using ImageJ (National Institutes of Health, USA). To enhance visual clarity, background removal was performed using a 50 pixel (0.162μm pixel^−1^) rolling ball radius, followed by contrast limited adaptive histogram equalization (CLAHE) with a block size of 19, histogram bins = 256, and a maximum slope = 6. Image channels were merged and flattened before being saved. Images were not subject to background subtract or CLAHE for any quantification purposes. Captured images were converted to a binary format, and the best fit ellipse was identified for each cell. The aspect ratio (AR) was defined as the ratio of the major to the minor axis of the ellipse. Statistical significance was tested using the non-parametric Mann-Whitney test (n = 387 cells on aligned substrate, and n = 139 cells on unaligned substrates), with p<0.05 being significant. Actin alignment was quantified using the OrientationJ plugin in ImageJ. In the text, data are described as mean ± standard deviation unless otherwise noted.

### Ultrathin porous parylene (UPP) membrane fabrication

Ultrathin porous parylene (UPP) membranes were fabricated using our previously published methods^26^. Briefly, six-inch diameter silicon wafers were coated with a water-soluble sacrificial layer of Micro-90 (International Products Corp, NJ, USA), followed by parylene-C deposition using an SCS Labcoter® 2 (Specialty Coating Systems, Indianapolis, IN). Parylene thickness measurement was performed using a Tencore P2 profilometer (KLA-Tencor, Milpitas, CA). Similar to our previous work^27,28^ standard photolithography was used to pattern a photoresist layer with hexagonally-packed circular pores with 8 μm diameter and 16 μm center-to-center spacing. This pattern was transferred to the parylene layer using inductively coupled plasma reactive ion etching (ICP-RIE) using a Trion ICP Etcher (Trion Technology, Inc., Tempe, AZ).

### Attachment of UPP membrane module

The UPP membrane was detached from the silicon wafer by immersing the wafer in DI water until the UPP floated away from the substrate (~1 minute). A membrane holding frame was fabricated using 1.5mm × 18mm PMMA (McMaster-Carr, California, USA) with a laser cut 2 mm × 2 mm region central was attached to the UPP using pressure-sensitive adhesive (MP468, MP467, 3M Company, Minnesota, USA). The outer edges of the PMMA frame were attached to the glass coverslip surrounding the exposed collagen gel using pressure-sensitive adhesive such that the UPP membrane made direct contact with the collagen.

### Confocal Reflectance Microscopy and Alignment Characterization

Collagen fiber structure was visualized using confocal reflectance microscopy (Leica TCS SP5 II) using a 488nm laser line with a 40x water objective and optical zoom of 1.75x. All settings were constant during the imaging sets. 2μm tall z-stacks were obtained in the center of the 60μm tall channels, each consisting of 13 images. The stacks were processed to an average projection plane before further processing. LOCI CT-FIRE (University of Wisconsin), a MATLAB based curvelet transform package, was used to quantify fiber directionality and anisotropy^29–31^. Anisotropy was quantified using the coefficient of anisotropy (CoA), which was calculated as the fraction of the fibers lying within ±15° of the mode, compared to the total number of fibers detected. Fiber directionality was reported as an angle with respect to the horizontal axis, as noted in the text.

## 3. CONTROLLING FIBER ANISOTROPY AND DIRECTIONALITY

Several recent papers have demonstrated that the extensional strain rate applied to a collagen solution can induce the formation of anisotropic collagen fibers. Paten and coworkers extended a needle away from the surface of a neutralized collagen droplet at a controlled rate such that the collagen was exposed to a constant extensional strain rate^32^. They reported that collagen fibers self-assembled in the direction of the needle motion and produced a collagen strand containing highly anisotropic fibers. Similarly, Lai and coworkers extruded acid-solubilized collagen from a needle moving at high velocity (340 mm s^−1^) onto a coverslip in a neutralizing bath of NaOH and PBS^5^. The resulting collagen contained anisotropic fibers in the direction of extrusion. Malladi and coworkers used a microfluidic flow-focusing technique to extrude a thin collagen sheet onto a mandrel with anisotropic fibers oriented in the direction of mandrel rotation^33^. Based on these results, we hypothesized that controlling the extensional strain rate applied to a self-assembling collagen solution within a microfluidic channel network would result in long-range (millimeter scale) domains of anisotropic fibers with fiber directionality defined by the flow path.

To test our hypothesis, we developed a non-uniform microchannel design, as shown in Figure 1A. The network consisted of five access ports, two in the ‘inlet region’ from 1-2, and three in the ‘expansion region’ from 3-5. The inlet and expansion regions were connected via a “funnel” with a 10:1 change in width from the funnel opening to the exit. With a constant input flow rate Q, and a varying cross-sectional area (i.e., channel width), the average fluid velocity, V, in the channel changed according to the relation V= QA^−1^. The network was designed to maintain a linearly changing velocity profile along the length of the funnel (Figure S1). The extensional strain rate was determined as the change in centerline velocity (ΔV) along the flow direction in the channel (Δx). A COMSOL flow simulation predicted an extensional strain rate in the fluid of 110 s^−1^ along the path 2-1 and 130 s^−1^ and in the funnel region. The fluid velocity dropped off non-linearly upon entering the expansion region (large increase in A), and the corresponding strain rate decreased from 130 s^−1^ to 30 s^−1^ over a 0.7 mm distance. The simulated flow velocity and streamlines are shown in figures 1C and 1D, respectively. An image of fluorescent beads following the flow streamlines at the entrance of the funnel is shown in Figure 1E.

To determine whether we could achieve fiber alignment by applying extensional strain to a collagen solution with our system, we first primed the microfluidic network with a neutralized solution of 2.5 mg ml^−1^ collagen (shown in dark blue in Figure 1A) at a flow rate of 4 *μ*l min^−1^ with all ports open. The pre-filling step was performed to limit bubble formation in the subsequent injection. As shown in figure 2A, Ports 3 and 4 were closed, and a neutralized 2.5 mg ml^−1^ collagen solution was injected into the microfluidic channel at a flow rate of 500 μl min^−1^ for 5 seconds with the flow directed along the paths 2-1 and 2-5 (shown in light blue). After injection, the devices were immediately transferred to a 37°C incubator for two hours to promote gelation. Confocal reflectance microscopy (CRM) images in the boxed yellow region show the collagen landscape over a 1.4 mm distance (Figure 2A). The degree of anisotropy (*i.e.*, coefficient of anisotropy, CoA) was quantified as the fraction of fibers within ±15° of the mode of the fiber angle histogram. In the boxed region, the CoA was maintained within 0.1 CoA unit (CoA = 0.60 to 0.69). We also observed that the fiber anisotropy was uniform throughout the gel (see supplemental video). In line with our hypothesis, the directionality of collagen fibers was controlled by the flow path. As shown in Figure 2A, the fiber directionality shifted from 0° near port 2 to 75° as the flow moved toward port 5, corresponding to a rotation of 53.6° mm^−1^. Thus, by defining a flow path (i.e., opening and closing access ports) in a microfluidic system with a non-uniform channel cross-sectional area, we could independently control fiber anisotropy and fiber directionality.

**Figure 2:**
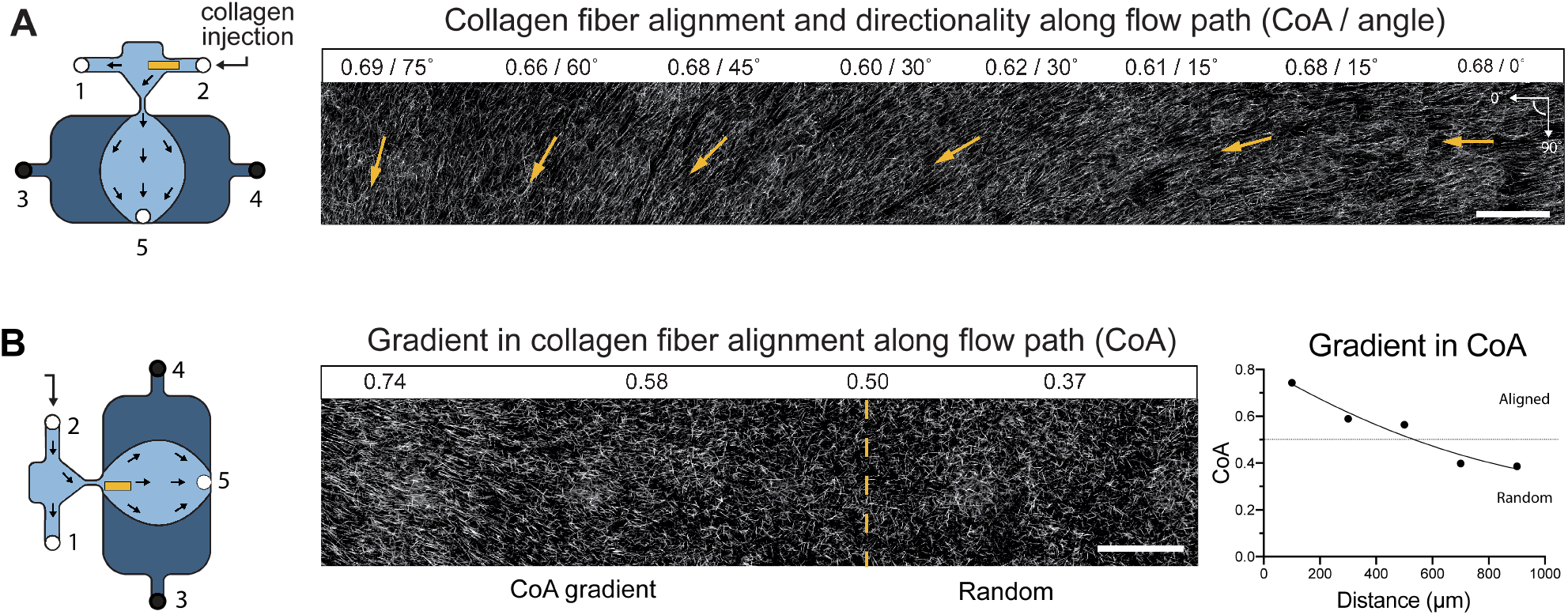
Confocal reflectance microscopy (CRM) images of COL1 fibers after injection and gel formation. The yellow box in each schematic identifies the imaging regions. (A) The stitched CRM image montage on the right and shows anisotropic fibers between ports 2 and 1, directed along the 0° axis at the inlet (port 2), and gradually turning towards the 90° axis into the funnel towards port 5. Arrows indicate the fiber directionality. A constant strain rate of 110 s^−1^ was predicted using a COMSOL flow simulation. (B) The stitched CRM image montage on the right shows that collagen anisotropy rapidly decreases over a 0.7 mm distance upon entering the expansion region. The gradient in CoA with a quadratic curve fit (R^2^ = 0.95). The slope in CoA from 0.74 to 0.5 was measured to be 0.0005 CoA units *μ*m^−1^. Scale bar = 100*μ*m in all images.

The dimensionless Weissenberg number (Wi) describes the elastic force to viscous force ratio in a non-Newtonian fluid; molecular stretching is favored when the elastic forces dominate (Wi ≥ 1). With a maximum predicted extensional strain rate of 130 s^−1^ in our channel, the Wi = 0.93 (detailed calculation in the supplement S2). While the Wi is slightly less than 1, our system has shear and extensional strain components that could contribute to the formation of anisotropic fibers^15,17,34^. Interestingly, anisotropic fibers were formed in our system even with a short injection process that lasted for ~ 5 seconds (total of 40uL injected along path 2-1 and 2-5). This observation can be partially explained because the application of strain has been shown to enhance the rate of polymer self-assembly by stretching protein molecules and increasing the number of nucleation sites. Thus, the fibers can preferentially nucleate along the flow direction (extension direction) and create anisotropic fibers after gelation^35^.

To further explore the relationship between the extensional strain rate and collagen CoA, we analyzed the fiber anisotropy at the end of the funnel as the collagen solution entered the expansion region. As shown in figure 1A and supplemental S1, our simulations predicted a rapid decrease in velocity resulting from the large increase in cross-sectional area. The corresponding extensional strain rate fell sharply from 130 s^−1^ to 30 s^−1^ over a 0.7 mm distance. As the strain rate decreased, the collagen fiber CoA reduced from 0.74 (anisotropic) to 0.5 (random orientation) over 500μm and could be described by a quadratic curve with an R^2^ = 0.95. To the best of our knowledge, this is the first demonstration of a controlled gradient in collagen fiber anisotropy within a 3D gel. Since the imposed extensional strain rate for a material is a function of the flow rate and channel geometry, we anticipate that CoA gradient characteristics such as the slope, profile, and boundary values can be tailored tuned by modifying the network design. Exploiting the relationship between extensional strain rate and CoA can provide new experimental capabilities to study how different cells migrate along domains of defined CoA as well as gradients in CoA within a 3D gel, with particular relevance to immune cell and cancer cell migration^36^.

We have also demonstrated that fiber directionality is a function of the defined flow path and can be controlled independently from fiber anisotropy. Combining well-defined characteristics of the collagen landscape in a single platform could enable future experiments that explore hierarchal interactions between fiber anisotropy and directionality that guide topographic cell alignment or migratory activities in a tissue-specific manner.

## 4. ESTABLISHING MULTI-MATERIAL INTERFACES

While controlling structural fiber properties is essential in creating an *in vitro* model, the material composition is an important design consideration. For example, the tumor microenvironment exhibits regions of dense hyaluronic acid (HA) and anisotropic COL1 type I collagen^37^, while more defined boundaries separating material domains are a hallmark of tissue barriers such as the transition from the stoma to the vasculature^38^. Although interfaces between materials can be achieved with sequential molding processes^39,40^, layer by layer ECM deposition^41,42^, viscous fingering^43^ and parallel microfluidic channels^44,45^, the microengineering of material interfaces while maintaining fiber anisotropy has not been widely explored. Here, we use our multi-port, two-stage injection process to control the fiber anisotropy and directionality, and exploit laminar flows of viscous solutions to define the interface between material domains. Since flow within our microfluidic network is in the laminar regime (maximum Reynolds number = 6), we hypothesized that our two-stage injection process would result in minimal mixing between gel solutions. To demonstrate the fabrication of material interfaces, we prefilled our microfluidic network with a 2.5 mg ml^−1^ COL1 solution, as described in the methods section. For the second stage of injection, a 2.5 mg ml^−1^ COL1 solution containing 4.8% HA (COL1-HA) was injected into the prefilled network along the path 2-1 and 2-5. Fluorescent beads were added to the COL1-HA solution to enable visualization of the interface. The samples were allowed to gel in a 37°C incubator before CRM imaging of fibers and fluorescence imaging of the beads. Figure 3A shows the schematic representation of the injection process, and the yellow box indicates the imaged region. Figure 3B shows the interface between COL1 and COL1/HA solutions identified by fluorescent beads. Figure 3C shows COL1-HA fibers aligned with a CoA of 0.68.

**Figure 3:**
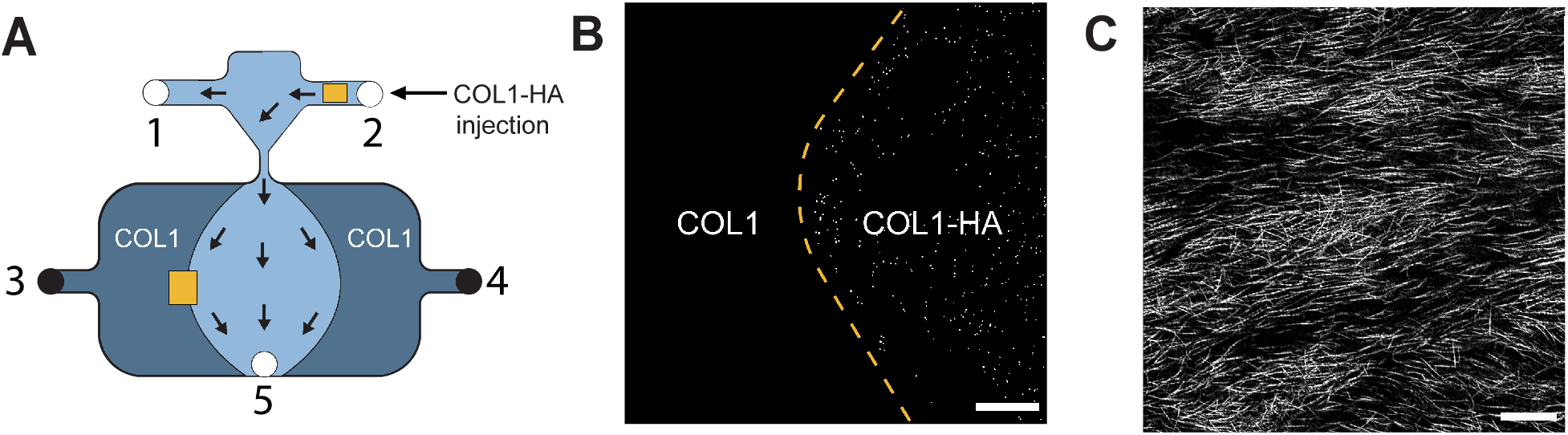
**(A)** Schematic showing the process of creating a multi-material interface using pure collagen (COL1) and collagen mixed with 4.8% hyaluronic acid (COL1-HA). (A) COL1 solution (dark blue) was injected from port 3 (pre-filling) with all ports open. Ports 3 and 4 were closed, and COL1-HA (light blue) was injected from port 2 along the path 2-1 and 2-5, as shown by the arrows. The yellow box in path 2-5 shows the region of interest imaged in panel B. **(B)** Fluorescent beads were incorporated into COL1-HA solution to visualize the interface between COL1 and COL1-HA after gelation. Scale bar = 500 *μ*m. **(C)** CRM image of anisotropic COL1-HA fibers (CoA = 0.67) in the region identified by the yellow box between ports 2-1 are shown. Scale bar 25*μ*m.

Thus, using a simple two-stage injection process along a user-defined flow path, we have combined two-material interfaces with anisotropic fibers in a single platform. The laminar flow and multi-port injection capabilities can be further leveraged to create more complex interfaces with structural definition and multi-material heterogeneity. Establishing independent control over the fiber anisotropy at a material interface could help shed light on questions related to cell migration, tissue morphogenesis, and early development.^46–48^

## 5. ACCESSING MICROENGINEERED COLLAGEN LANDSCAPES

Conventional microfluidic systems consist of channels that are permanently bonded to a substrate, with fluidic ports being the only point of access to the interior of the system. The introduction of a 3D gel into the channels effectively blocks these initial access ports^38,49,50^. Cells can be introduced to the system in two common ways: first, by pre-mixing them into the gel solution before injection^38^, or second, through the use of peripheral access channels separated from the main channel with small constrictions^49^. The use of adjacent channels to introduce cells is not always an option in large channels due to limited access to the gel’s interior. Including cells in the gel precursor solution is convenient but limits the extent to which fiber properties (*e.g.*, diameter, length, and density) can be controlled by tuning solution pH, ionic strength, composition, and polymerization temperature, because those variations can cause cell damage^51–53^. To overcome these practical limitations, we demonstrate a versatile method for accessing the 3D microengineered collagen landscapes after formation.

To address the access problem intrinsic to sealed microfluidic systems, we developed a method where the PDMS guiding channel could be lifted off after collagen gelation. The PDMS channels were passivated with BSA after fabrication to limit the attachment of collagen. Glass coverslips were functionalized with poly(octadecene maleic-alt-anhydride) (POMA) to promote covalent attachment of the collagen to the glass^39,54^. After collagen microengineering and gelation, the PDMS guiding channel was gently lifted off with tweezers and removed to expose the gel, as shown in Figure 4A. To assess how fiber anisotropy was affected by the lift-off process, we randomly selected 50 regions on the landscape and imaged them with CRM before and after channel removal. As shown in Figure 4 (B-C), the average change in CoA was found to be 0.0005 ± 0.05 (Figure 4C). CRM imaging also showed that the difference in the height of the collagen gel as a result of lift-off was minimal, with an average change of 1.4*μ*m ± 0.7*μ*m (Figure S3). Since the landscape properties and gel height was minimally impacted by channel lift-off, our technique represents a promising method to create well-defined collagen landscapes and provide easy access to the interior of the gels. Open access allows standard cell seeding and culture, fluorescent labeling, and RNA extraction protocols to be followed and, thus, improves the usability of the platform outside of engineering-focused laboratories.

**Figure 4:**
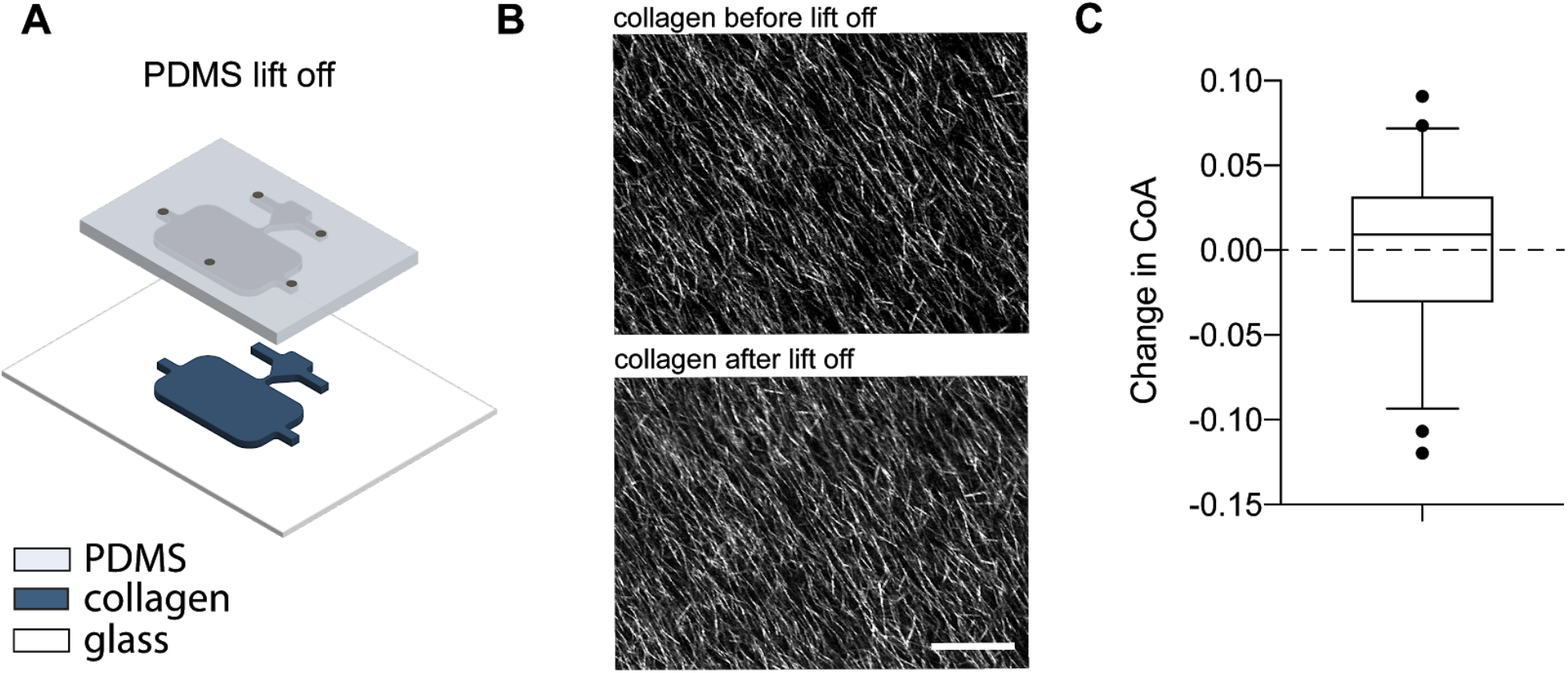
**(A)** Schematic representation of the PDMS channel lift-off process. Removing the PDMS channel after gel formation allowed direct access to the underlying collagen substrate (dark blue). **(B)** CRM images showing collagen alignment in a region of interest before and after lift-off. Scale 25 *μ*m. **(C)** Box and whisker plot showing the change in CoA across 50 randomly selected regions after before and after lift-off, with a mean change of 0.0005 ± 0.05 CoA units. Whiskers show 5-95 percentiles.

## 6. RESPONSE OF MDA-MB-231 AND HUVECS TO ALIGNED COLLAGEN

Next, we sought to confirm that cells would recognize and respond to the exposed collagen fiber landscapes after lift-off. MDA-MB-231 cells are an invasive triple-negative breast cancer cell line commonly used as a model to study matrix interactions. Here, we seeded them as single cells rather than aggregates to explore how they responded to the collagen fibers. An acrylic media reservoir was positioned around the collagen gels, and MDA-MB-231 cells were seeded on the gel and fixed after 24 hours. As seen in figure 5A, individual cells in the region between ports 2-1 aligned with along the collagen landscape following a path from ports 2-5. The alignment and direction of cells in the yellow boxed region were observed to be similar to the collagen landscape shown in Figure 2A. As shown in Figure 5B, cells interacting with aligned collagen fibers displayed a more elongated morphology as quantified with the cell aspect ratio (AR) when compared to cells on unaligned gels. The average AR of aligned cells was found to be 3.44 ± 2.4 compared to 2.13 ± 1.3 on randomly oriented fibers. These results show that cancer cells followed the anisotropy and directionality in our microengineered gels. Moreover, since the direction of collagen fibers is directed by channel design, cell orientation and alignment can be achieved by modifying the channel design.

**Figure 5:**
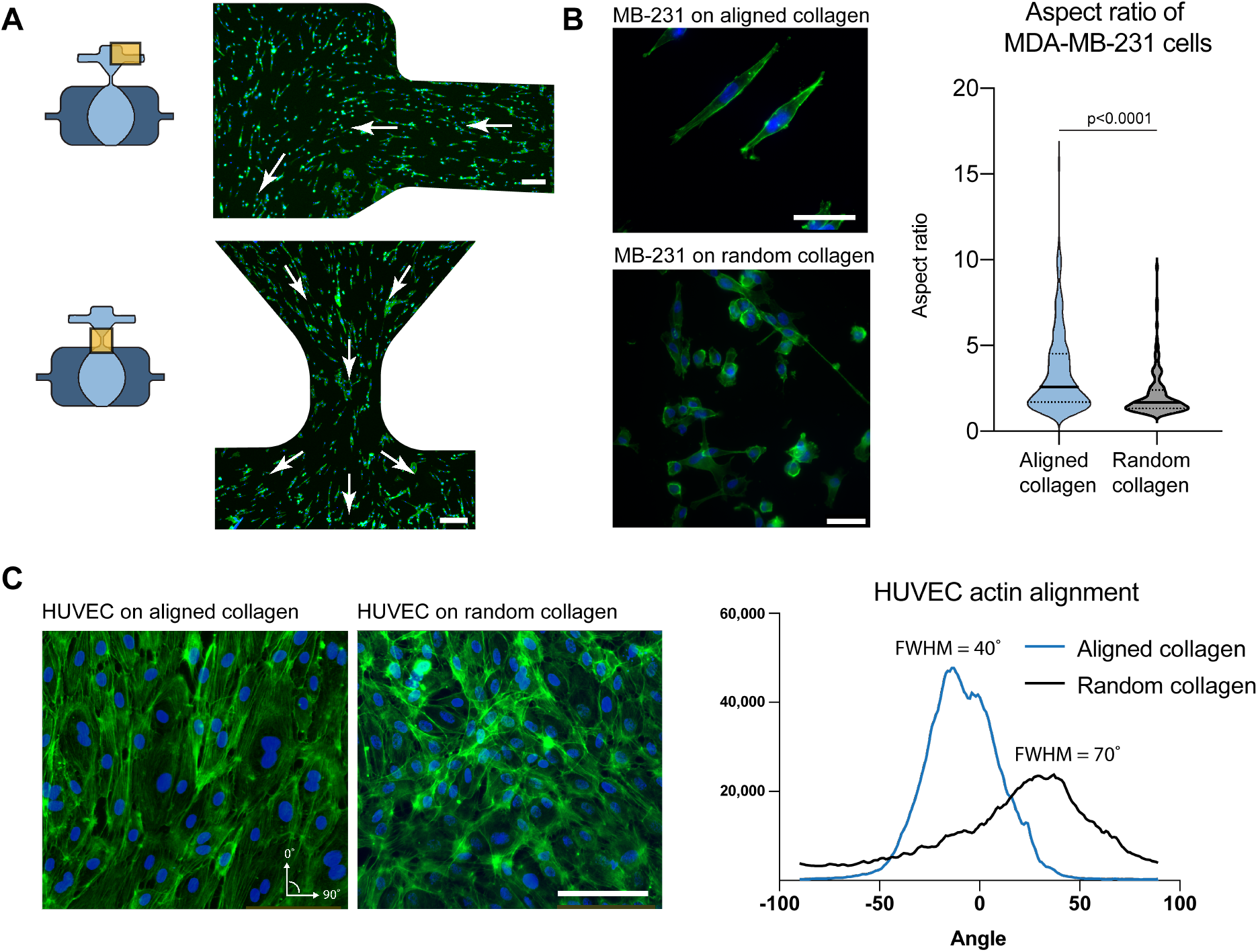
(A) MDA-MB-231 cells were fixed and stained to identify actin (green) and nuclei (blue) 24 hours after seeding on aligned collagen landscapes. Yellow boxes in the schematic show the imaged regions. Cells were observed to be elongated on aligned fibers and followed the fiber direction, indicated by the white arrows. (B) Cells on aligned collagen displayed a more elongated morphology as compared to cells on randomly oriented collagen gels. The plot compares the cytoskeletal aspect ratio of cells on aligned collagen and random collagen. Aligned cells had a mean aspect ratio of 3.44 ± 2.4 compared to a mean of 2.13 ± 1.3 for randomly oriented cells. (C) HUVECs on the collagen were stained for actin (green) and nuclei (blue) after 24 hours. Cells on anisotropic gels displayed aligned actin fibers. The histograms show the distribution of HUVEC actin fibers on aligned collagen and random collagen, with FWHM values of 40° and 70°, respectively.

Next, we seeded human umbilical vein endothelial cells (HUVECs) to test whether a sheet of cells would respond to the collagen landscape. Figure 5C shows images of HUVECs on aligned and random collagen after 24 hours, stained for actin (green), and nuclei (blue). HUVECs on both gels can be seen forming a confluent sheet, with the actin fibers aligning in response to the anisotropic collagen fibers (FWHM = 40°) while cells on random collagen gels did not show preferential alignment (FWHM = 70°).

The above results demonstrate that single cells and cell monolayers both respond to the collagen features in our gels after the lift-off process. The sequential workflow of microengineering the gel and then seeding cells onto the exposed surface addresses concerns about matching gel self-assembly and alignment-promoting factors (e.g., pH, ionic strength, temperature, and extensional strain) with cell compatible conditions. After fabrication, the landscape can be rapidly equilibrated with media to ensure the gel is cell compatible. This is relevant in primary cell populations that are particularly sensitive to their local environment. In our platform, the channel design and flow properties can be tailored to control the collagen landscape characteristics (CoA, gradients in CoA, interfaces, and fiber direction), and the resulting landscape properties guide cell behaviors. In this way, we leverage the precise fluidic control and manipulation capabilities hallmark to microfluidic systems with the streamlined and established cell culture and analysis protocols used in conventional cell biology settings. Direct access to the gel substrate also enables the addition of secondary modules to improve functionality, as discussed in the next section.

## 7. DEMONSTRATION OF MODULAR CAPABILITIES USING AN ULTRATHIN POROUS PARYLENE MEMBRANE

The ability to lift off the PDMS channel is beneficial because we can expand the capabilities of our platform through the addition of application-specific modules. Modular approaches to microfluidic systems have been demonstrated previously using open microfluidics^55,56^, magnetically sealed devices^57,58^, and LEGO-type blocks^59^. One of the key advantages of using modular microfluidic systems is the ability to discretize an experiment into specific components and add or remove them as required during the experimental workflow.

To demonstrate the modular capability of our platform, we introduced an ultrathin porous parylene (UPP) membrane as a mask to define the degree of interaction between cells and the underlying collagen. As shown in Figure 6A, the UPP was attached to a laser cut PMMA reservoir and placed on the exposed collagen gel after channel lift-off. The UPP was fabricated in-house, with a thickness of 863 nm, with a pore size of 8μm and 23% porosity^24^. Figure 6B shows a CRM image where aligned collagen fibers are visible under the UPP (dark regions are pores in the membrane). The aligned collagen fibers were seen making contact with the membrane’s pores after the application of the UPP module. MDA-MB-231 cells were seeded onto the membrane surface and stained for actin (green) and nuclei (blue) after 24 hours. Figure 6C shows that cells could grow on the UPP and make contact with the underlying aligned collagen. However, actin fiber alignment was not observed, possibly due to decreased cell-collagen interactions.

**Figure 6:**
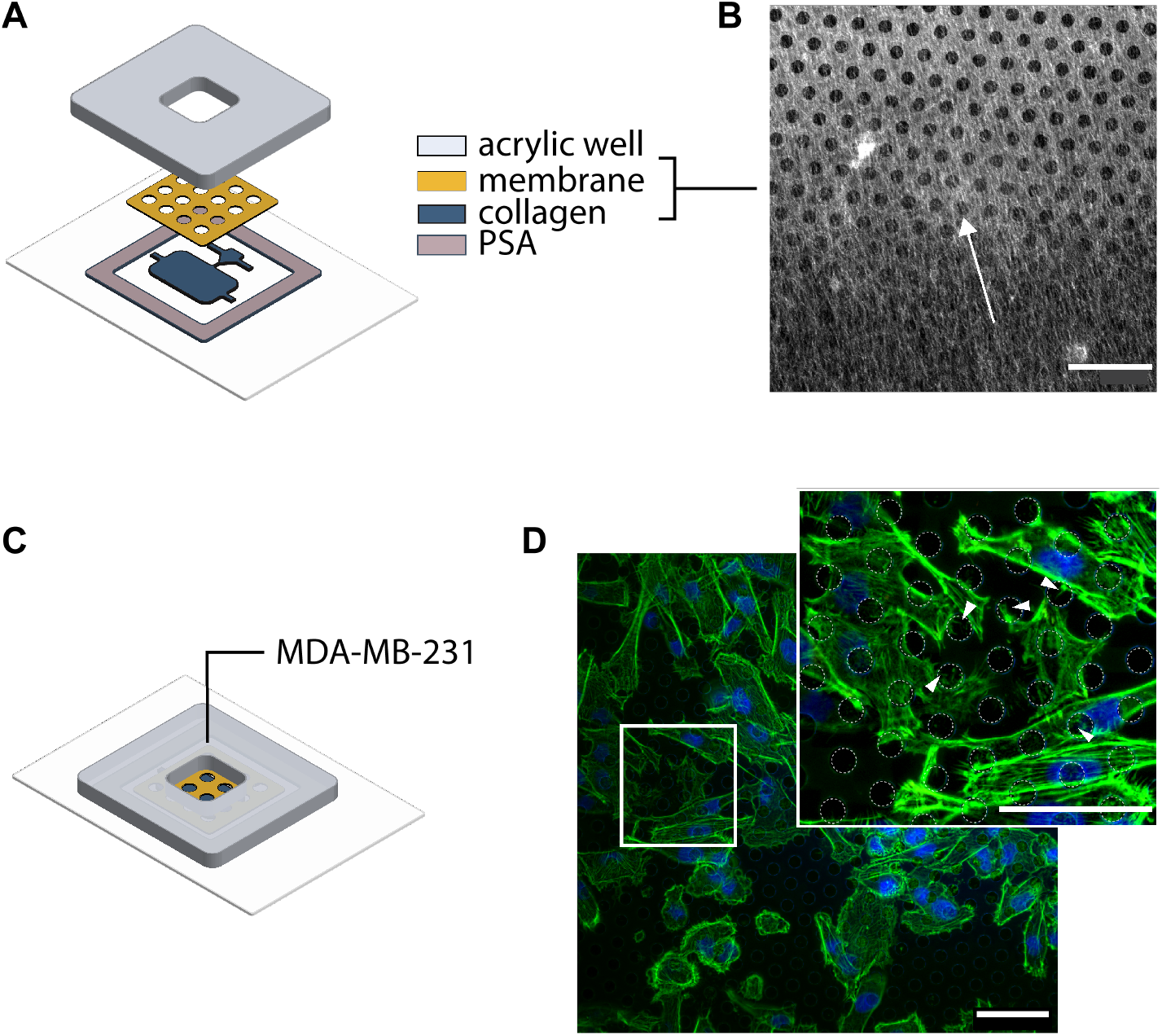
(A) Schematic of the membrane placed on the exposed collagen landscape using pressure-sensitive adhesive and PMMA well for holding cell media. (B) CRM micrograph of the membrane and aligned collagen fibers in contact with the membrane. The image is captured from the underside of the membrane. The arrow indicated collagen fiber directionality. (C) Schematic showing the process of seeding cells on UPP. (D) MDA-MB-231 cells were seeded on the membrane and stained after 24 hours for actin and DAPI to visualize cell morphology. The image inset shows distinct filopodia-like protrusions formed by cells over the membrane pores, suggesting that cells can probe the underlying ECM through pores, indicated by white arrows. (Scale bar = 50*μ*m).

Interestingly, distinct filopodia-like protrusions (white arrows) were extended by the cells in Figure 6D, inset, suggesting that the cells were probing the collagen under the pores. These results indicate that a similar platform could be used to carry out further studies to optimize the pore size and spacing to understand how much cell-ECM interaction is required to induce cell alignment. The use of membrane modules also have importance in modeling barrier tissue functions such as studying the effects of ECM alignment on leukocyte transmigration^60^, or improving the adhesion of cells on a membrane under an applied shear stress^61^

## 6. CONCLUSIONS

In this work, we have demonstrated a platform that combines microfluidic fiber alignment capabilities with fiber directionality control traditionally associated with extrusion-based methods. We showed that the fiber directionality could be controlled by changing the fluid flow path (i.e., opening and closing of access ports) and fiber anisotropy could be defined by changing the extensional strain rate using expanding and contracting channel geometries. In addition, both the flow path and strain rate can be controlled as a function of the channel geometry; therefore, modification of collagen landscapes can be achieved by simply changing the channel design and experimental flow parameters. We also showed that we could achieve gradients in fiber anisotropy and exploited laminar flows to create defined interfaces between materials while maintaining anisotropy. We further demonstrated a lift-off process to simplify experimental workflows and showed that cells responded to the microengineered collagen landscape, and confirmed that a UPP membrane module could be added to the system to explore cell sensing behaviors. We anticipate that additional modules such as flow channels, soluble factor reservoirs, or sensors can be added to tailor the experimental platform to fit a broad range of applications. We believe that the versatile capabilities provided by our system will be of interest to researchers looking to recapitulate tissue-specific microenvironments *in vitro*.

## Supporting information

Z-stack of aligned collagen gel

## ABBREVIATIONS

CRM: Confocal Reflectance Microscopy
HA: hyaluronic acid
COL1: type 1 collagen

## ASSOCIATED CONTENT

### Supporting Information

The following files are available free of charge.

Video showing aligned collagen through the gel thickness (movie 1, gif)

Supplemental calculations and channel dimensions.

## Author Contributions

The manuscript was written through contributions of all authors. All authors have given approval to the final version of the manuscript.

## ACKNOWLEDGMENT

The authors acknowledge assistance from Dr. Hyla Sweet in the Confocal Microscopy Lab, and Dr. Christopher Lewis and Mr. Tom D. Allston from the Materials Characterization Laboratory for assistance with rheometry.

