## Supplemental Figures for "Microengineered three-dimensional collagen fiber landscapes with independently tunable anisotropy and directionality"

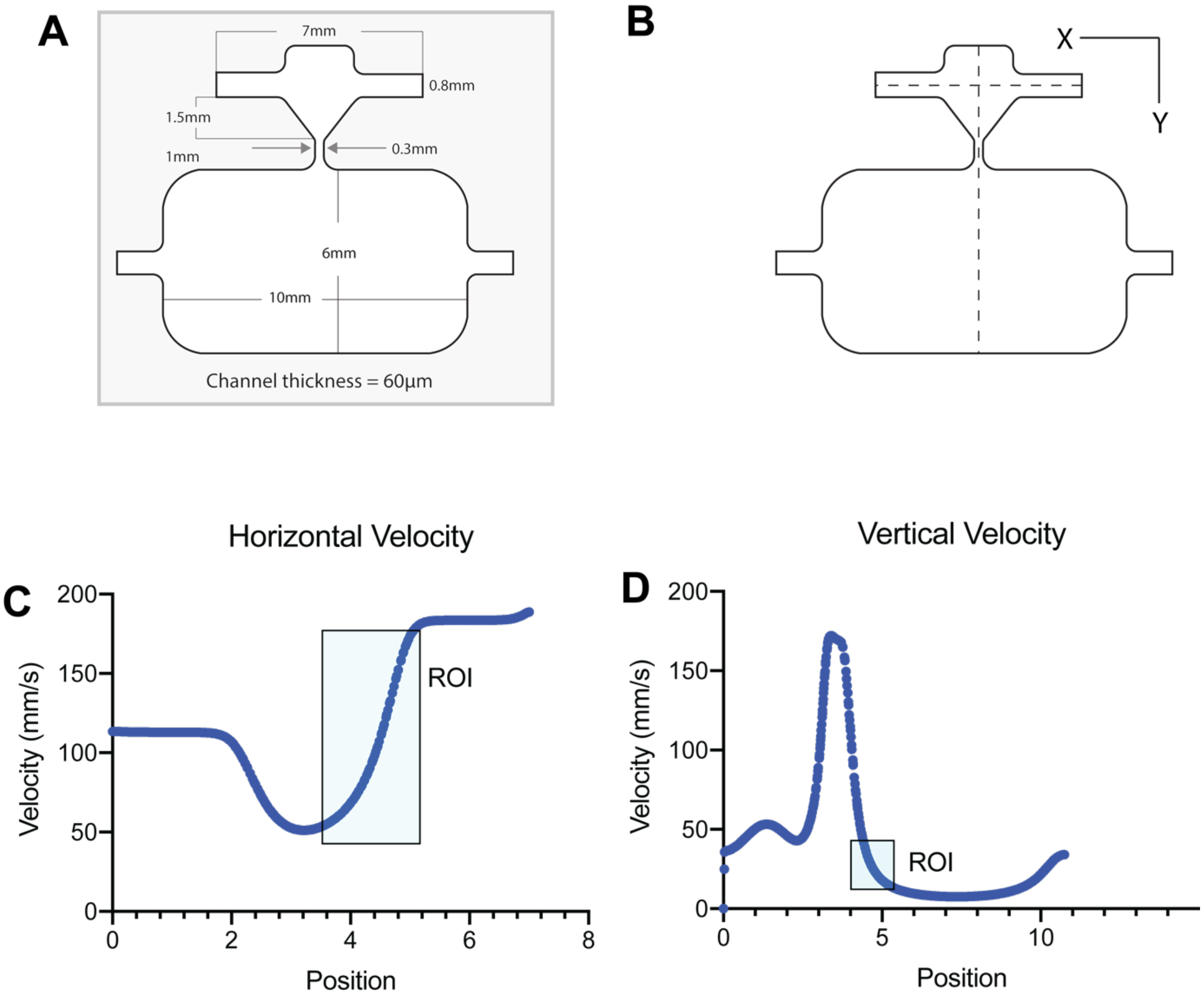


Figure S1: (A) Channel dimensions (B) Schematic showing the line plots along which velocity was measured (C,D) Line plots of velocities showing the regions of interest used to calculate the extensional strain rate

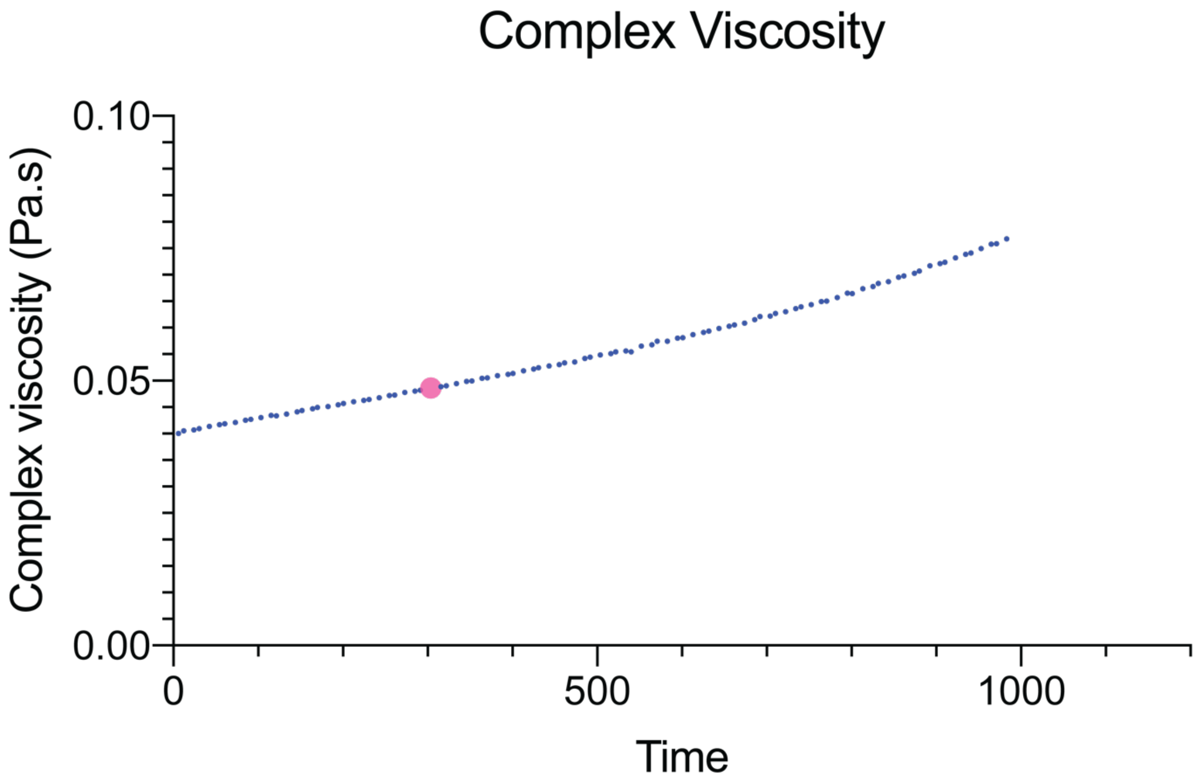


Figure S2: Plot of complex viscosity as measured using rheometry at 21˚C. Each point represents average of 5 independent experiments. The average value after 300s was calculated to be 0.00498 Pa.s (49.8cP)

Weissenberg Number (Wi) Calculation:

Relaxation time ($\tau$)= 144*viscosity (μs) [1]
= 144*49.8 = 7171.2 μs = 0.0071 s

Max strain rate in channel ($\gamma$) = 130 $s^{-1}$

Wi max = $\gamma\tau$ = 130*0.0071 = 0.92

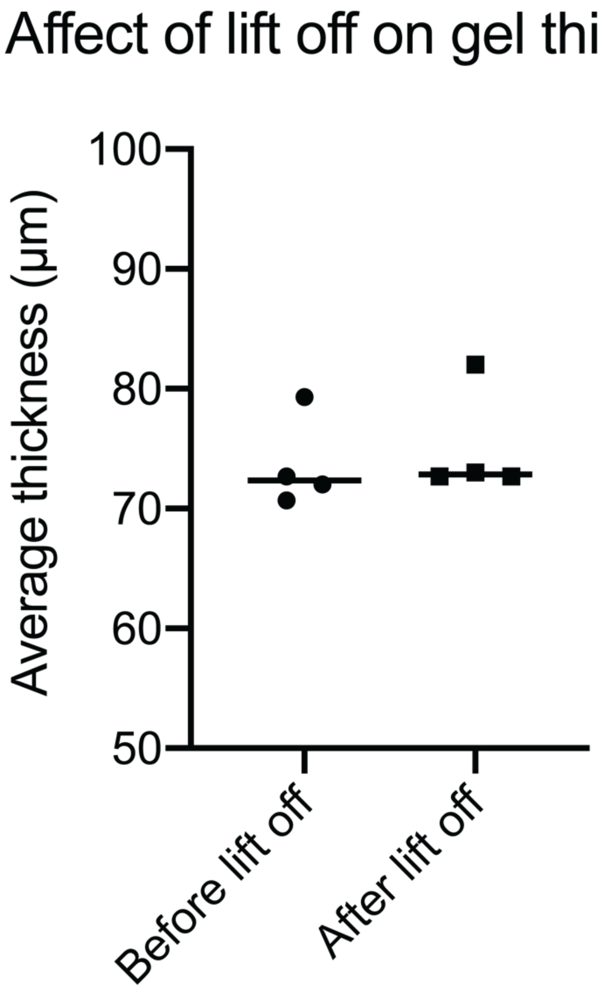


Figure S3: Change in the thickness of collagen gel after PDMS channel lift off. Each data points represents the average of 3 points on a sample. Average change in height after channel lift off was determined to be 1.4µm

(1) Yamakawa, H. Viscoelastic Properties of Straight Cylindrical Macromolecules in Dilute Solution. *Macromolecules* **1975**, *8* (3), 339–342. https://doi.org/10.1021/ma60045a019.
